# Duckweed roots are dispensable and are on a trajectory toward vestigiality

**DOI:** 10.1101/2022.01.05.475062

**Authors:** Alex Ware, Dylan H Jones, Paulina Flis, Kellie Smith, Britta Kümpers, Levi Yant, Jonathan A Atkinson, Darren M Wells, Anthony Bishopp

## Abstract

Duckweeds are morphologically simplified, free floating aquatic monocots comprising both rooted and rootless genera. This has led to the idea that roots in these species may be vestigial, but empirical evidence supporting this is lacking. Here we show that duckweed roots are no longer required for their ancestral role of nutrient uptake. Comparative analyses of nearly all rooted duckweed species revealed a highly reduced anatomy, with greater simplification in the more recently diverged genus *Lemna*. A series of root excision experiments demonstrated that roots are dispensable for normal growth in *Spirodela polyrhiza* and *Lemna minor*. Furthermore, ionomic analyses of fronds in these two species showed little difference in the elemental composition of plants in rooted versus root-excised samples. In comparison, another free-floating member of the Araceae, *Pistia stratiotes*, which colonized the aquatic environment independently of duckweeds, has retained a more complex root anatomy. Whilst *Pistia* roots were not absolutely required for growth, their removal inhibited plant growth and resulted in a broad change in the mineral profile of aerial tissues. Collectively, these observations suggest that duckweeds and *Pistia* may be different stages along a trajectory towards root vestigialization Given this, along with the striking diversity of root phenotypes, culminating in total loss in the most derived species, we propose that duckweed roots are a powerful system with which to understand organ loss and vestigiality.

**One sentence summary:** Through their adaption to the aquatic environment, duckweed roots have progressively become structurally reduced making them an ideal plant model with which to study vestigiality.

## Introduction

Evolution has shaped the body plans of all organisms into the myriad of diverse forms we see today. While evolution is commonly envisioned as constantly generating novel forms, things sometimes go the other way: occasionally, entire structures or traits are lost, becoming vestigial. This can result in radical shifts in body plan and life-history strategy and is a key evolutionary process driving structural innovation. Based on earlier definitions (Prout, 1964, Fong et al., 1995, Müller, 2002), vestigiality can be broadly defined as the retention, through evolution, of genetically determined structures that have lost some or all of their ancestral function.

Vestigiality is phylogenetically widespread in plants (Knobloch, 1951). Examples include loss of entire organs, such as floral organs in *Penstemon sp*., oil glands in *Ceratandra* flowers, leaf reduction in *Equisetum*, and non-functional roots in dodder seedlings, and are often concurrent with atypical, innovative body plans or unusual life history strategies (Walker-Larsen and Harder, 2001; Sherman et al., 2008; Steiner, 1998). To date, reports exploring vestigiality in plants are largely descriptive. Progress into understanding the molecular and evolutionary processes which drive organ loss in plants has therefore been limited.

The most advanced work driving our understanding of the molecular control of vestigiality comes from outside the plant kingdom. Perhaps the most detailed work has been done on the blind cavefish, *Astyanax mexicanus*, where the mechanisms underpinning eye-loss have been well described. Comparisons between blind and sighted cavefish has revealed that lens apoptosis is mediated by expansion of the expression domain of *sonic hedgehog A* and *B* (*shhA* and *sshB*), which negatively regulate the homeobox gene *pax6*, itself a key regulator of eye development in vertebrates (Yamamoto et al., 2004). It is clear from work in *Astyanax* that leveraging the presence and absence of organs in closely related species is crucial to gaining an understanding of vestigiality at a molecular level. We propose that root loss in duckweeds represents a powerful untapped model for understanding organ loss in plants due to the existence of closely related rooted and rootless species. Recent development of genetic tools (Yang et al., 2018; Vu et al., 2020; reviewed in Acosta et al., 2021) has enabled exploration of molecular networks in duckweeds. However, any study of vestigiality first requires a detailed understanding of how the organ in question functions. As roots are still retained in many duckweed species, we need clarity on duckweed root function to frame the evolutionary context of this model. Within the literature there are several observations regarding the function of duckweed roots; however there is no single study bringing together multiple lines of empirical evidence supporting their vestigiality.

Duckweeds are highly morphologically reduced free-floating angiosperms lacking many of the key organs common in flowering plants, such as clearly defined stems and leaves. The plant body is reduced to a flattened frond or thallus. They comprise five genera divided into two subgroups, Lemnoideae (*Spirodela, Landoltia* and *Lemna*) and Wolffioideae (*Wolffia* and *Wolffiela*). Within these genera, there is an evolutionary trajectory in root number consistent with root vestigialization: the earliest-diverging duckweed genera (*Spirodela* and *Landoltia*) possess multiple roots, later diverging ones a single root (*Lemna*), and the most recently diverging lineages possess no roots at all (*Wollfia* and *Wolffiela*) (Tippery and Les, 2020, Figure 1).

**Figure 1.**
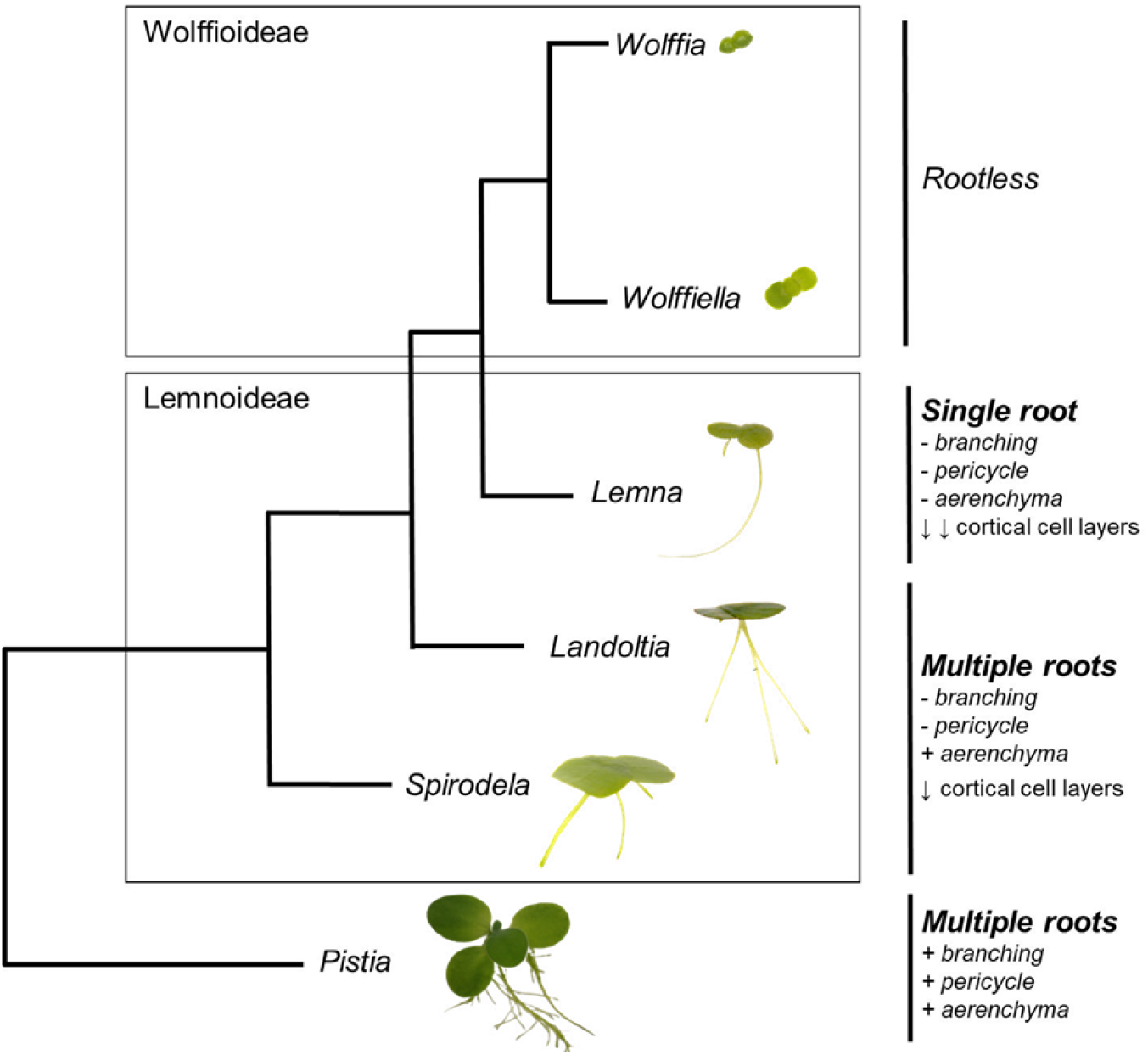
Representative phylogeny of the duckweed genera and *Pistia* highlighting the progressive loss of roots of roots and loss of individual root traits as genera diverge (indicated by + and -; arrows next to cortical cell layers indicate the progressive reduction in layer number as the genera diverge) (after Tippery and Les, 2020). Representative images (not to scale) of species from each genera are shown for illustrative purposes.

Duckweed roots are adventitious and neither branch nor form root hairs (Landolt, 1998). Previous studies have performed detailed investigations into root anatomy in individual species, reporting high levels of structural reduction. *Spirodela polyrhiza* roots have a stele comprising of one xylem cell, two sieve elements and between five and six phloem parenchyma cells (Kim, 2007). These are enclosed by a single layer of endodermis, three distinct cortical cell layers and between 38-45 epidermal cells (Kim, 2007). A similar pattern is reported for *Lemna minor* (Echlin, 1981). Although there have been other studies of root anatomy (eg. Hegelmaier, 1868), we currently miss a systematic understanding of root anatomy across the three root-bearing genera.

Vestigialization not only affects anatomy, but also function. It does not imply that organs should possess no function, only that the salient function is lost. Here, we define the salient function of roots as organs with which to acquire water and nutrients. Various lines of evidence have been presented to support the view that duckweed roots have at most a limited role in nutrient uptake. Hegelmaier (1868) noted that in their natural habitat, individuals of *Lemna gibba* without roots occur. Gorham (1941) concluded that nutrients were taken up via fronds and not roots, as coating the underside of fronds with a hydrophobic wax reduced the division rate of fronds and caused root elongation, whilst coating the upper surface did neither. Muhonen and colleagues (1983) also noted that *Spirodela polyrhiza* grew without roots. Whilst these studies suggest that roots may not be required for growth, they do not rule out that duckweed roots still play some role in resource capture. Indeed, it has been observed that both roots and fronds can assimilate nitrogen in both *Lemna minor* and *Landoltia punctata* (Cedergreen and Madsen, 2002; Fang et al., 2007).

The above presents an incomplete picture of vestigiality in duckweed roots. To address this, we conducted a survey of duckweed root anatomy across almost all the rooted duckweed species. We examine to what extent changes in anatomy are consistent with roots being vestigial, and if additional structural reduction accompanies the reduction in root number between genera. We then investigated root function in two species by looking at growth and uptake of 13 elements in plants with and without roots excised. By comparing duckweeds with the related free-floating macrophyte *Pistia stratiotes* we present a scenario in which both anatomical complexity and the role of the root in foraging for nutrients has been progressively lost in duckweeds.

## Results

### Duckweed root anatomy is highly reduced

Previous reports of duckweed root anatomy focused on just a few species and have not directly compared these with relatives. Without outgroups, it is impossible to determine if there is a trajectory towards structural reduction in duckweeds. Previous phylogenetic studies have included *Pistia stratiotes* as an outgroup as another aquatic member of the *Araceae* (Les et al., 2002). *Pistia* and duckweeds share several morphological and ecological similarities as free-floating macrophytes but represent independent aroid lineages (Stockey et al., 1997, Wilde et al., 2005). Indeed, both fossil evidence (Stockey et al., 1997, Wilde et al., 2005) and phylogenetic analyses (Friis et al., 2004) suggest that duckweeds and *Pistia* independently colonized aquatic habitats, with fossils attributable to the duckweeds being much older than those attributable to *Pistia* (Cabrera et al., 2008). Thus, *Pistia* provides a useful model for understanding the highly reduced structure in duckweeds as its form resembles ancient fossil duckweeds such as *Limnobiophyllum*.

We surveyed macroscopic root structure for 20 duckweed lines, representing 13 species, with all *Spirodela* and *Landoltia* species represented and 10 of the 13 *Lemna* species, alongside *Pistia stratiotes*. Most species were represented by multiple accessions. In no instances were lateral roots or root hairs observed in duckweeds, in line with previous observations (Landolt, 1986). *Pistia* had a considerably larger and more complex root system with lateral roots. 11 out of 12 duckweeds have a mean root diameter between 120-200 µm, with only *Lemna yungensis* falling outside of this range of means, possessing a mean diameter of 256 µm. No duckweed species possessed a maximum root diameter close to the 325 µm that we observed in *Pistia*. We counted an average of 212 total cells in *Pistia* cross-sections, while duckweeds display mean total cells values of 28-81 cells. *Spirodela* sp. displayed mean cross section cell numbers ranging from 40 to 81 cells. *Lemna* species typically displayed fewer cells in cross-section than *Spirodela*; mean values for all species are between 28 and 45 cells, apart from *Lemna yungensis*, which displays 73. Morphological analysis of root patterning revealed a highly reduced anatomy common to all duckweed species. This consisted of a 3-5 of cortical cell layers and a highly reduced vasculature (Figure 2A). All the duckweed species possessed a single central xylem, typically surrounded by a small number (7-10) of what appear to be phloem parenchyma cells, although this identity has never been explicitly defined. *Pistia*, conversely, has multiple xylem files and considerably more phloem cells. It also has a discernible pericycle, which was absent in all duckweeds surveyed here (Figure 2A, Figure 3, Supplementary Figure 2).

**Figure 2.**
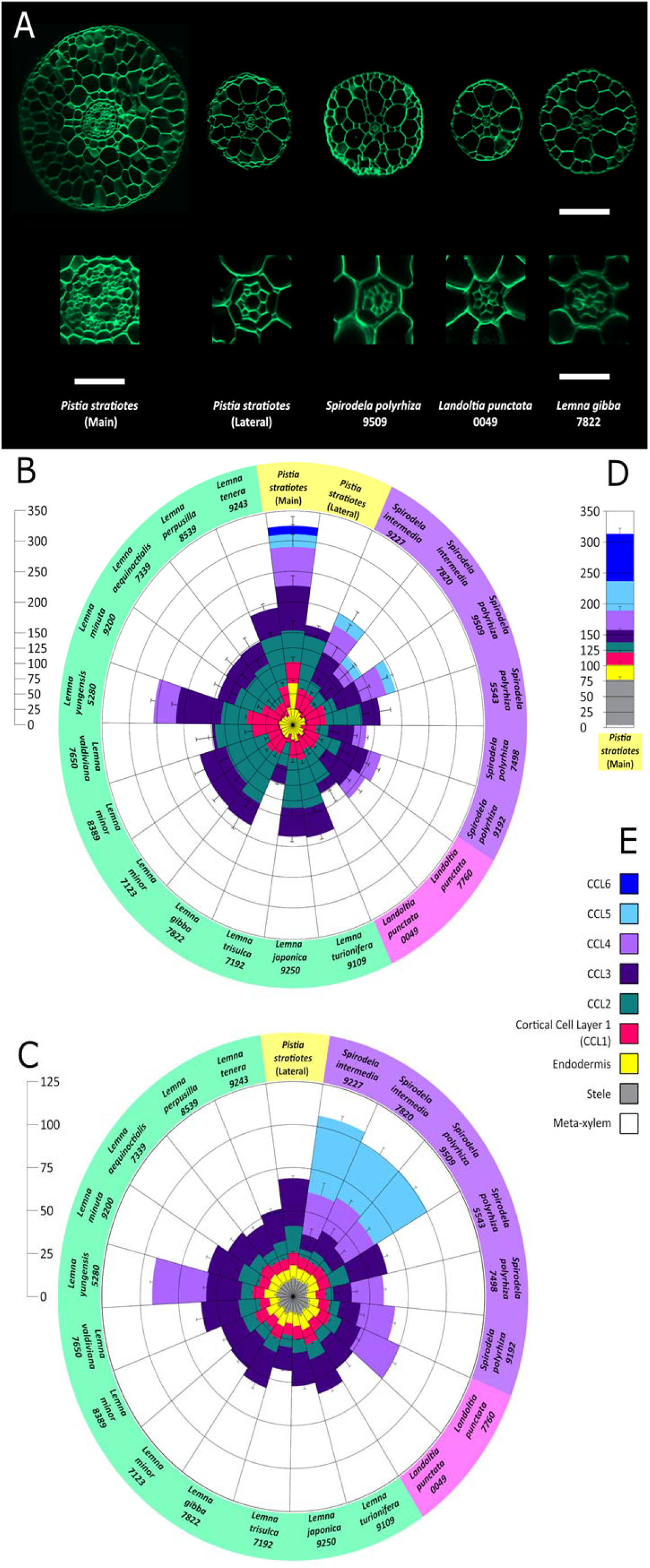
Comparison of root anatomical traits across almost all extant duckweeds reveals a highly reduced anatomy. A) Representative images of root sections from species representing each duckweed genera and mainand lateral roots of *Pistia stratiotes*. Images were obtained via fresh tissue sectioning and confocal imaging. Scale bar = 50 µM for entire roots; 10 µM for vasculature close-up. B) Rose diagram displaying the width of each cell layer (µm) for roots of 20 duckweed lines encompassing 13 species, denoted at the outside of the circle. C) Rose diagram displaying the number of cells in cell layer for roots of the aforementioned lines, denoted at the outside of the circle, with *P. stratiotes* main roots (D) in a separate bar chart for ease of resolution. Background colour underlying the species labels represents genera; yellow represents *Pistia*, purple *Spirodela*, pink *Landoltia*, green *Lemna*. E) Colour coded key to the different cell layers displayed on the rose diagrams. CCL stands for cortical cell layer. *n* = 10 root sections derived from different plants, except for *Pistia stratiotes* (main) and *L. trisulca* where *n* = 5 root sections derived from individual plants.

**Figure 3.**
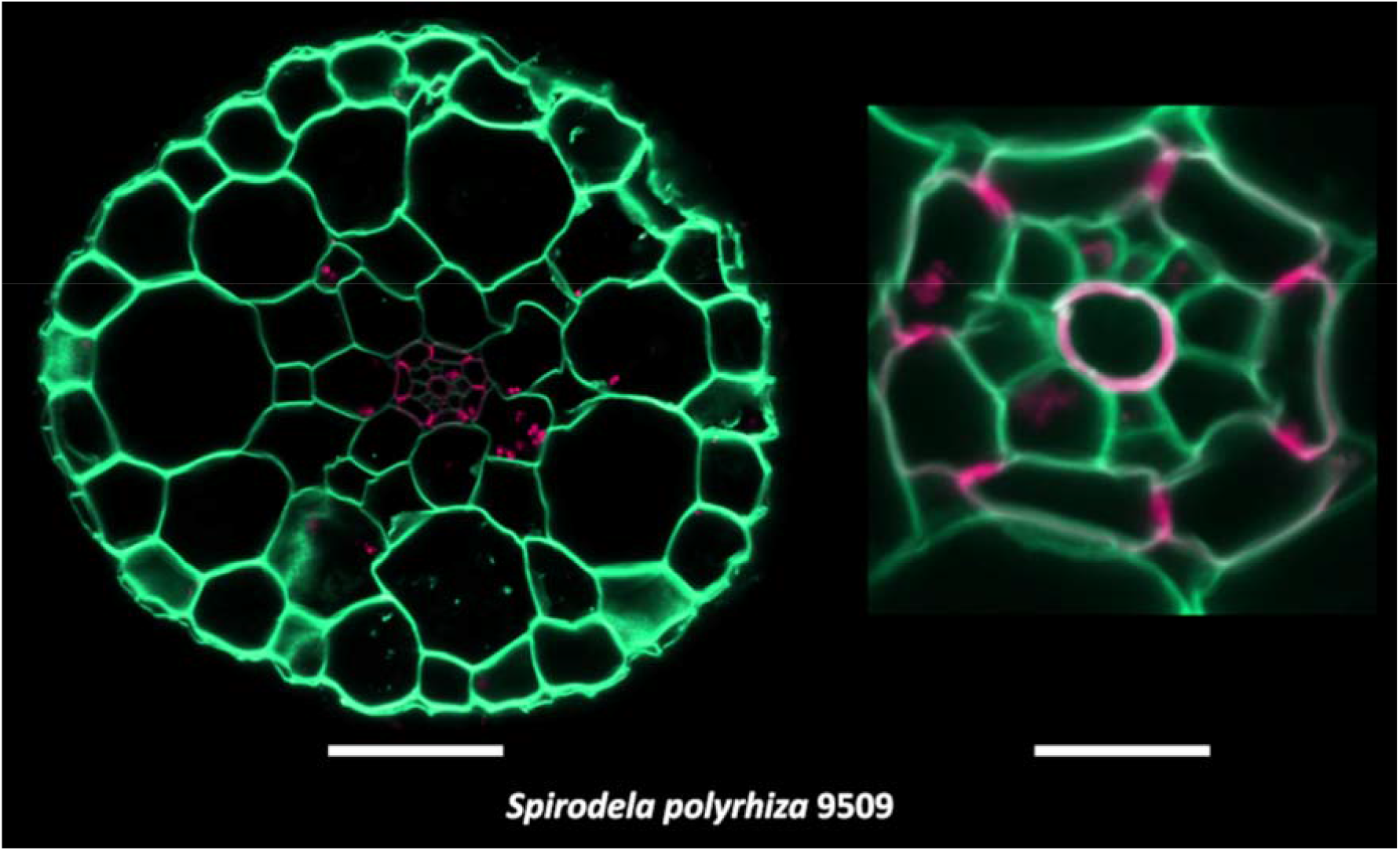
Basic fuchsin staining of duckweed vasculature highlights lignification in the endodermis and central xylem. Entire root section and accompanying close up of the vasculature of *Spirodela polyrhiza* 9509 with cell wall staining (calcofluor white; green) and lignin staining (0.01% Basic Fuchsin; magenta). Scale bar = 50 µM for entire roots; 10 µM for vasculature close-up.

We observed a trend in the reduction of number of cortical cell layers (CCLs) from the earlier diverging *Spirodela* (3 of 6 accessions display 5 CCLs) to the later diverging *Lemna* (3 CCLs in 11 out of 12 accessions) (Figure 2B). This trend is not reflected in the root diameter (Figure 2C). Several duckweed species have large extracellular air spaces within the cortex, similar to the schizogenous aerenchyma found in many other aquatic plants (Jung et al., 2008). This feature appears more frequently in *Spirodela* (5 out of 6 lines), in 1 out of 2 of the *Landoltia* lines, and in only 2 out of 12 closely related *Lemna* lines that are currently proposed to represent a single species (*yungensis valdiviana*) (Bog et al., 2020) (Figure 2A, Supplementary Figure 1).

Compared with *Pistia*, the cell number and size of the stele and endodermis is uniformly low across all duckweed species. The total number of cells enclosed by and including the endodermis is remarkably invariable across duckweed species, with all duckweed species falling within the range of 16-18 cells, compared to approximately 100 in *Pistia*. The diameter of the endodermis is slightly more variable than cell number, with the mean for all species within a range of 15-28 µm, with no clear pattern between genera. The fact that duckweeds consistently showed reduced cell size and number within the stele suggests reduced importance for transport within the root, consistent with vestigiality.

We quantified a number of parameters relating to number and size of each cell type in the root and conducted a principal coordinates analysis to survey the general trends in this anatomical dataset (Figure 4). Each point represents the data captured from a root section of a separate individual (Figure 4A), and the 19 variables are shown in (Figure 4B). The PCoA displays 5 distinct clusters, consistent with phylogenetic groupings. All *Lemna* species are retained in a single cluster (3), apart from *Lemna yungensis*, which forms a distinct unique cluster outside of the larger *Lemna* cluster containing this species alone (4). *Spirodela intermedia* occurs in two distinct clusters (1 and 2), neither of which contain any *Lemna* individuals. The majority of *Spirodela intermedia* individuals cluster together, in a group also consisting of a small number of *Spirodela polyrhiza* samples (cluster 2). *Spirodela polyrhiza* is distributed more broadly and located within clusters 1, 2 and 3, with the majority of individuals falling into cluster 2, which contains only *Spirodela* and *Landoltia* species. A small number of *Spirodela polyhriza* samples also fall into the *Lemna* cluster. *Landoltia* primarily co-occurs with *Spirodela polyrhiza* and *intermedia* in cluster 2, and a few individuals occur in the *Lemna* cluster. *Pistia* main roots group distant from all duckweeds driving the main axis, PC1. Interestingly, all *Pistia* lateral roots fall within the *Lemna* cluster. Given that the duckweed genera broadly cluster within their own groups, and that we see a reduction in root complexity (CCLs & aerenchyma) from *Spirodela to Lemna*, we propose that root anatomy is progressively reduced in more recently derived duckweed lineages.

**Figure 4.**
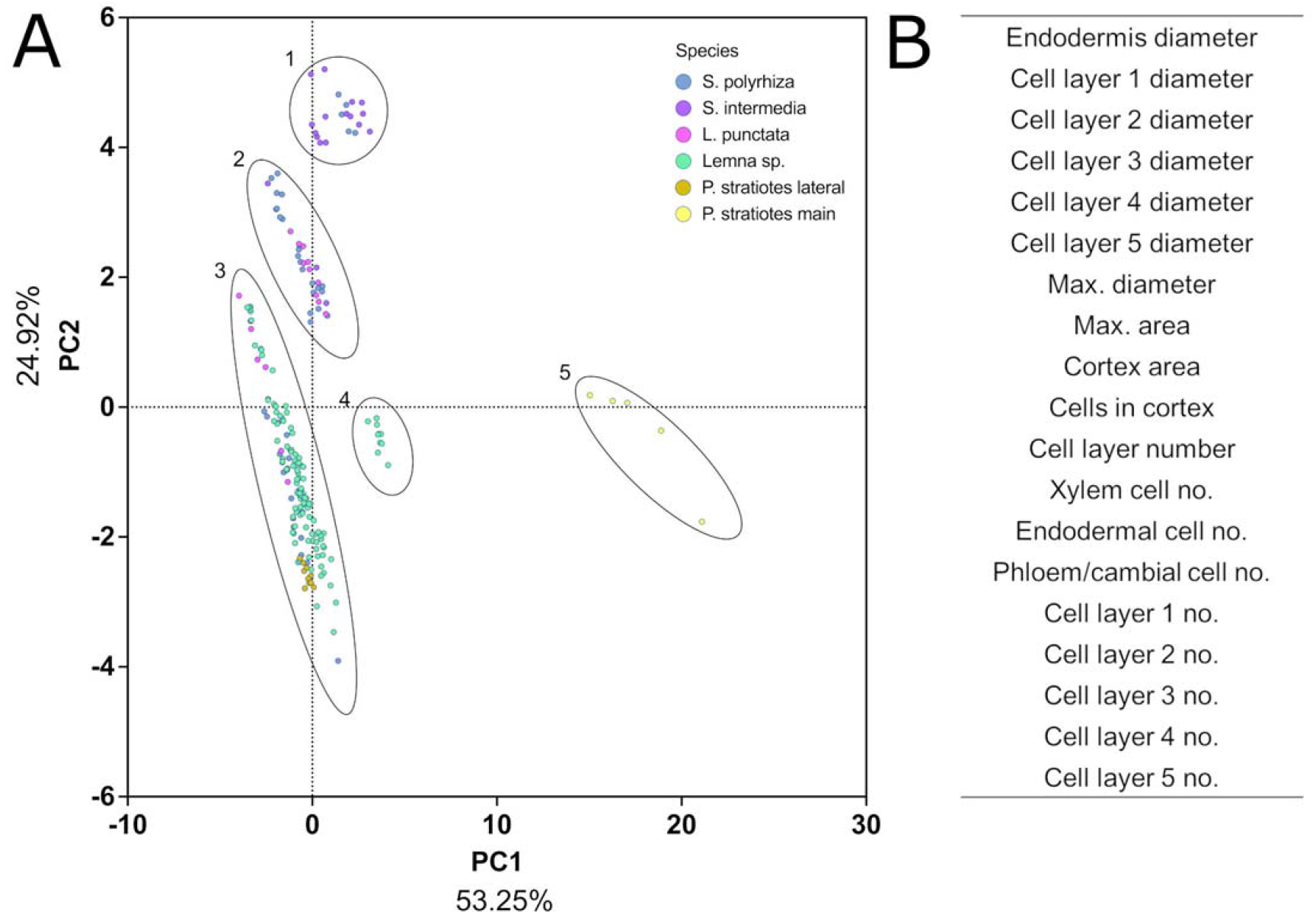
Principal coordinates analysis of duckweed anatomical traits highlights interspecies differences and a gradient of reducing root anatomical complexity. A) PCoA based on 21 components, with 210 rows, derived from an anatomical analysis of fresh root sections from 20 duckweed lines, encompassing 13 species, and main and lateral roots of *Pistia stratiotes*. Clusters have been manually highlighted and numbered for ease of further discussion. Percentage of variance explained by each PC is indicated on the relevant axis. B) Summary of the 19 variables used to generate the PCoA in A.

### Continuous root removal does not reduce duckweed growth, but does reduce growth in *Pistia stratiotes*

We hypothesised that a reduction in root complexity would be reflected by reduced requirement of roots for plant growth. To test this hypothesis, we conducted root removal experiments and compared the growth rate response to root removal in two representative duckweed species, *Lemna minor* and *Spirodela polyrhiza*, alongside *Pistia stratiotes*. Root removal was conducted daily for a period of 11 days to minimise growth of new root material. Growth (as frond or aerial tissue area) was measured daily, normalised as a percent of the initial area value (Figure 5). During the growth series, we observed an approximate 12 fold increase in frond area for *Lemna minor*, a 10 fold increase for *Spirodela polyrhiza*, and an 8 fold increase in area for *Pistia stratiotes* for individuals in control samples where roots were intact (Figure 5). For *Spirodela polyrhiza* (Figure 5A) we saw no significant difference in growth for rooted versus root-excised samples. In *Lemna minor*, the only significant differences in growth arose on the final three days of the growth series, where plants with their roots removed displayed enhanced growth (Figure 5B). In contrast, root removal markedly reduced the growth rate of *Pistia stratiotes* (Figure 5C). These results indicate that duckweed roots are not required to sustain growth in laboratory conditions. These results also suggest that the root is not an essential means of water absorption in duckweed.

**Figure 5.**
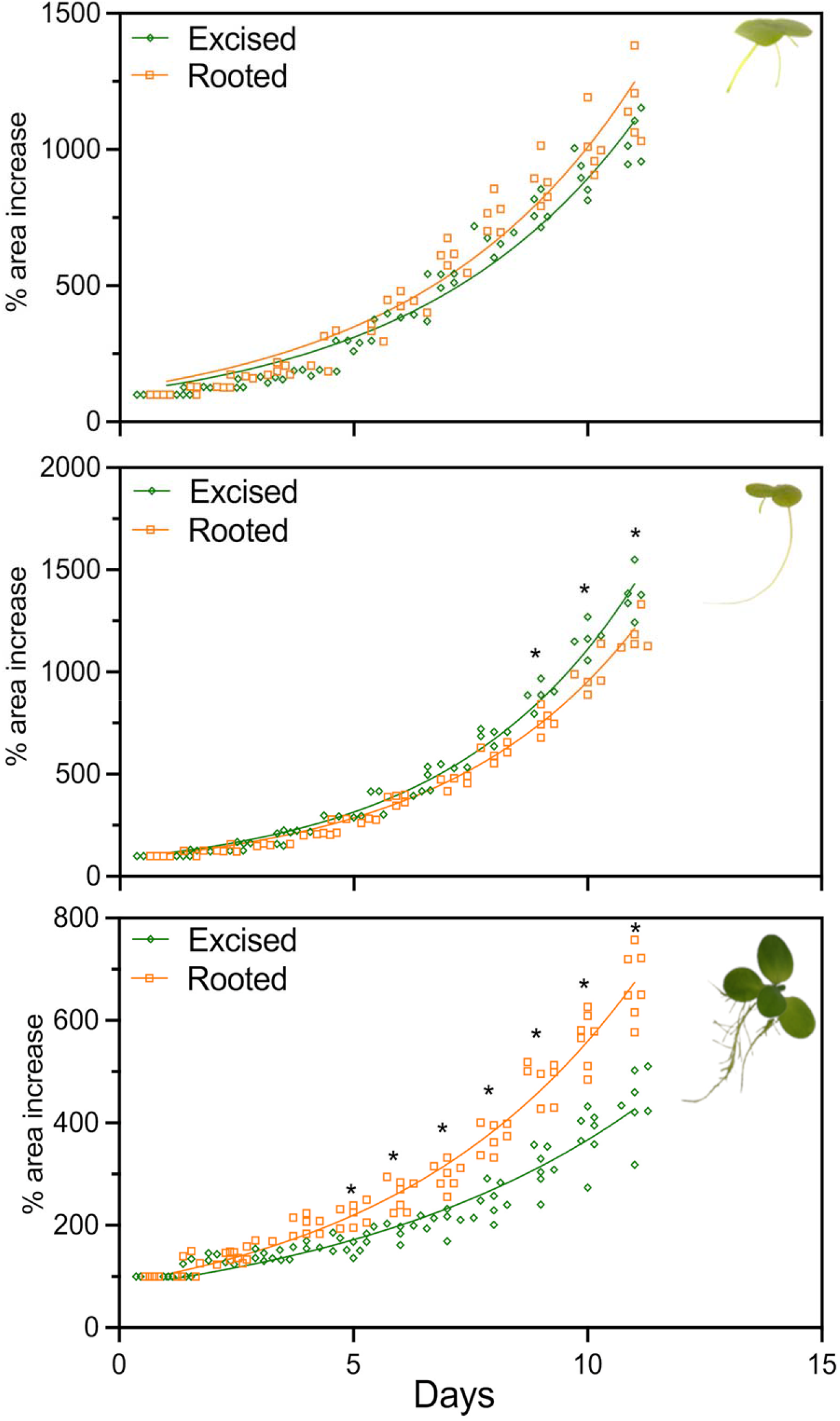
Growth of the duckweeds *Spirodela polyrhiza* and *Lemna minor* is not impacted by continual root removal, unlike the aroid *Pistia stratiotes*. Plants were subjected to continuous root removal and growth compared to untreated controls. Growth was measured as area of fronds (or aerial tissues for *Pistia*), derived from daily imaging from beneath, and plotted as a percentage increase relative to the initial (day 1) area value. Lines show the best fit of an exponential growth curve. A) *Spirodela polyrhiza*; B) *Lemna minor*; C) *Pistia stratiotes. n* = 5 flasks, each initially seeded with 3 colonies for duckweeds; *n* = 7 flasks, each initially seeded with 1 plant for *Pistia*. Asterisks show statistically significant differences as assessed by two-way repeated measures ANOVA followed by Sidak’s multiple comparisons. Lines show the best fit of an exponential (Malthusian) growth curve.

### Root removal does not impair the ability of *Lemna minor* or *Spirodela polyrhiza* to absorb macro- and micronutrients, but does impact nutrient uptake in *Pistia stratiotes*

The growth rate assay established that rooted versus root-excised duckweeds grew in a similar manner, but root removal impeded the growth of *Pistia stratoites*. We reasoned that if roots were required for the uptake of specific elements, assays in which we measure specific elements would be more sensitive than a crude measurement of growth in detecting the extent to which roots are still required for their ancestral function. To investigate this, we subjected the fronds and aerial tissues of *Pistia* generated by the previous experiment to an ionomic analysis. A total of 16 elements were successfully detected in these species. Whilst some rare elements such as Li and Cd were detected, we only considered 13 elements present in our growth media B, Na, Mg, P, S, K, Ca, Fe, Mn, Co, Cu, Zn and Mo (supplementary figure 2). As our analysis was run under atmospheric conditions, we were unable to measure the levels of N.

Root removal in duckweed made little change to the overall accumulation of nutrients in duckweed (Figure 6A, B). Between *Lemna minor* and *Spirodela polyrhiza*, there were five instances where root removal significantly altered elemental concentration in the frond. In three instances, root removal resulted in a significantly up-regulated accumulation of certain elements: we saw increased Ca concentration in the fronds of both *L. minor* and S. *polyrhiza*, and increased Fe, Zn and Mn in *S. polyrhiza* alone. Root removal resulted in a reduction in concentration of B and Cu in *L. minor* alone (Figure 6A, B, Supplementary Figure 2). In contrast, the impact of root removal on the ionomic composition of *Pistia* was considerably greater, with P, S, K, Fe, Mn and Zn all being significantly reduced (Figure 6B, Supplementary Figure 2). Together, these data suggest that whilst roots are no longer required for growth and nutrient uptake in duckweeds, *Pistia* roots still play an important role in growth and nutrient acquisition. However, given that they are not absolutely required for growth, it may be that *Pistia* is *en route* to root vestigiality, albeit at a less advanced stage than the duckweeds.

**Figure 6.**
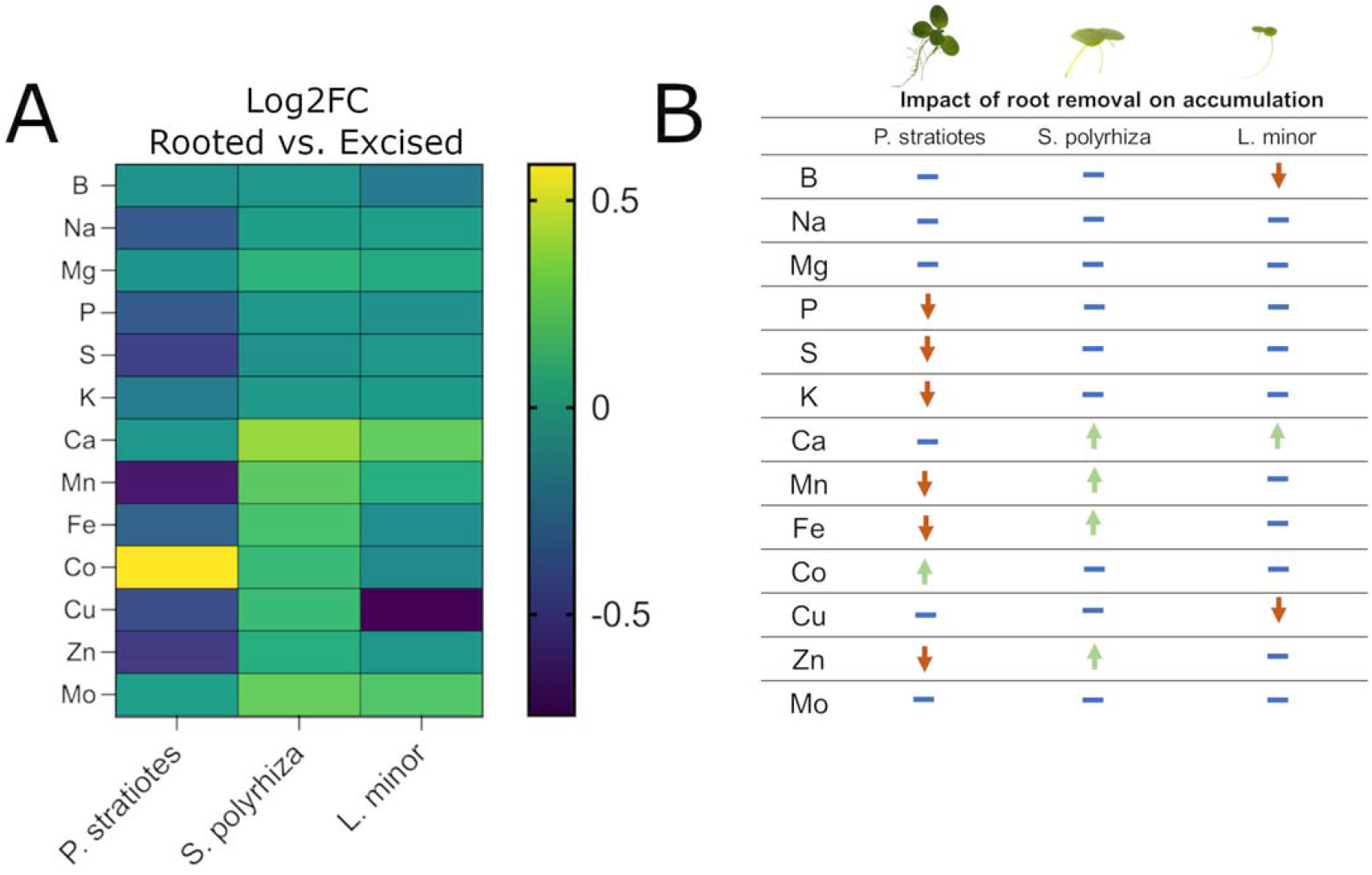
Continuous root removal has a limited effect on element accumulation on the duckweeds *Spirodela polyrhiza* and *Lemna minor* but reduces the accumulation of a number of elements in the aroid *Pistia stratiotes*. A) Heatmap showing the log2 fold change of rooted versus rooted elements for each species. B) Table synthesising the data generated in A) indicating whether root removal results in statistically significant increased accumulation (green upwards arrow), decreased accumulation (red downwards arrow), or no significant change (blue hyphen). Significance (P <0.05) was determined by one-way ANOVA followed by Sidak’s multiple comparisons test. *n* = 5 flasks, each initially seeded with 3 colonies for duckweeds; *n* = 7 flasks, each initially seeded with 1 plant for *Pistia*.

## Discussion

Here we sought to better resolve whether the duckweed root may be a vestigial organ, with the aim of clarifying if duckweeds may serve as a helpful model for understanding the molecular mechanisms underpinning organ loss. For an organ to be considered vestigial, it must have lost its salient function. Typically, such organs undergo accompanying reductions in size and complexity. Defining salient functions for an individual organ is challenging. However, it is clear that for almost all angiosperms, a primary function of roots is to supply water and nutrients to the growing plant, sustaining growth of the aboveground tissues (Boyce, 2005). We therefore examined the anatomy, as well as water and nutrient uptake ability, of duckweed roots to better ascertain the position of each species group along a trajectory towards vestigiality, culminating in root loss in the most recently evolved *Wolffia* and *Wolffiella* (Fig. 1).

We began by surveying the anatomy of a global collection of specimens including almost all rooted duckweeds, allowing us to observe if a) the reduced anatomy in duckweeds is consistent between species and genera and b) if any trends in root reduction are present at the anatomical level. This built upon previous reports looking into a handful of species (An et al., 2019; Landolt, 1986; Melaragno and Walsh, 1976), expanding it considerably to encompass almost all rooted species of duckweeds. We compared duckweed root morphology with the sister *Pistia stratoites*, which is believed to have undergone an independent and more recent invasion of the aquatic environment. Our findings revealed that duckweed roots are consistently reduced in both size (diameter) and morphological complexity compared with *Pistia*, consistent with the idea that they are no longer required for active nutrient transport (Figure 2A, Supplementary Figure 2).

As well as the macroscopic reduction in root system complexity - multiple roots per frond to single root per frond - in *Spirodela* and *Landoltia* versus *Lemna*, we also leveraged our anatomical data to question whether root anatomical complexity reduces concurrently with root number. We observed a reduction in both the number of cortical cell layers and the presence of aerenchyma between *Spirodela* spp. and *Lemna* spp. The apparent decrease in complexity between *Spirodela* spp. and *Lemna* spp. supports a model in which traits associated with root complexity have been progressively lost in duckweeds as novel species have formed, accompanying the reduction in root number. In comparison, *Pistia* plants may be less far along this trajectory towards root vestigialization. A PCoA encompassing all root anatomical traits measured further confirmed these observations. Virtually all individuals of the genera *Spirodela* and *Landoltia* sit in two distinct clusters based on their root anatomy, separated from *Lemna* individuals, which exist almost exclusively in a single cluster, matching their monophyletic origin. This correlation with phylogenetic groupings further supports the concept that root anatomy has evolved to become further reduced in *Lemna*. We feel root loss in duckweed presents a unique opportunity for deepening our understanding of vestigiality. In other models of organ loss, such as cavefish, evolution has produced a more binary range of traits (i.e. sighted versus unsighted fish). In comparison the duckweed root offers a greater spectrum of phenotypes in terms of both root number and anatomy, providing a rich pool of germplasm within which we can explore networks controlling discrete aspects of root development.

The anatomy of the duckweed root is also highly similar to that of lateral roots in *Pistia*. This cellular arrangement is similar to that of fine lateral roots of other monocot species (Watanabe et al., 2020). When root anatomical trait values are mapped onto a PCoA, *Pistia* lateral roots sit in a cluster which is primarily composed of *Lemna spp*. It is feasible that this cellular arrangement seen in *Lemna* represents or is approaching an anatomical ‘minimum’ without which it would not be possible to form a root.

If duckweed roots are vestigial, they should not only have reduced complexity but will have lost some or all of their salient function. We showed that whilst *Pistia* roots had a positive and significant effect on leaf growth, growth of duckweed fronds was largely unaffected in rooted versus rootless samples, implying that roots are dispensable for providing nutrients and water for growth. Growth data alone do not provide a full picture of capacity for nutrient transport. We therefore leveraged an ionomics platform that permitted a survey of the elemental landscape of duckweed fronds when grown without a root, which we compared with *Pistia stratiotes*. We considered a broad suite of nutrients including every element present in our growth media, except nitrogen. We did not see major shifts in the elemental composition of the fronds of either duckweed species when subjected to continuous root removal. Strikingly, no elements included in our analysis (0 out of 13) exhibited reduced accumulation in *Spirodela polyrhiza* grown without roots, and only 2 out of 13, B and Cu, did in *Lemna minor*. Conversely, in *Pistia stratiotes*, 6 of the 13 elements quantified exhibited reduced accumulation in shoot tissues as a consequence of root removal, including elements critical for growth with well-established root-mediated uptake mechanisms such as P and K. Together, this clearly evidences the dispensability of roots in duckweeds for nutrient uptake. A surprising result in both *Spirodela* and *Lemna* was the increase in certain nutrients following root excision. This included Ca, Fe, Zn, and Mn, with Ca being consistently elevated. A potential hypothesis is that duckweed roots could be repurposed for the storage or sequestration of nutrients. Raphides (calcium oxalate crystals) are present in *Lemna minor* and have been shown to localise within roots (Franceschi 1987, 1989).

Considering the definitions of vestigiality by both Prout and Muller, we feel that these data clarify that duckweed roots are indeed vestigial, and to varying degrees across the group, opening the door to their utilisation as models for understanding this vestigiality. This gradient also poses a key question. If duckweed roots are vestigial, why are they maintained in some species? Whilst some vestigial structures may be non-functional, others may have gained novel functions as a consequence of reduced constraint (i.e., exaptation), whilst other structures may be in an intermediate state whereby the transition to vestigiality is incomplete (Walker-Larsen and Harder, 2001). It is therefore possible that relaxed selection pressure has permitted duckweed roots to become neofunctionalised to perform novel roles. It has been suggested that duckweed roots may function as organs of stability (Landolt, 1986) or aid dispersal by adhering to animals (Cross, 2017).

In conclusion, these results support a model of progressive vestigiality of roots across the duckweeds. Broadly it points to a duckweed root that is both anatomically simplified and dispensable for the salient functions of water or nutrient uptake. However, we acknowledge that our experiments do not completely rule out a role for root in nutrient uptake under, for example, limiting conditions or in natural habitats, replete with companion species and competitors. However, these results lay a foundation for the use of duckweed roots as a model system for further investigation into the molecular and evolutionary processes underlying vestigiality in plants.

## Methods

### Duckweed growth and culture

All duckweed stocks employed in this experiment were obtained from the Landolt collection, ETH Zurich (http://www.duckweed.ch), except for Spirodela polyrhiza lines 9509 and 7948 which were provided by Klaus Appenroth, Friedrich Schiller University, Jena. Four-digit numerical codes following species names refer to their Landolt accession number. Stocks were maintained on liquid N-media or SH-media (Appenroth et al., 1996) at 120 µmol m^-2^ s^-1^ light and 16/8h light cycle in a Conviron growth chamber, set to 22°C with 70% RH. *Pistia* stratiotes was obtained from JAM Aquatics, Wrexham, UK.

### Root cross section anatomy

Plants were grown in 250 ml conical flasks containing 150 ml of liquid N-media in the same conditions as stocks. Flasks were inoculated with 5-10 colonies from the stock collections and grown for 2-6 weeks. Plants selected possessed roots of average or greater length, and fronds of average or greater area based on visual appraisal.

Vibratome sectioning of duckweed roots was conducted as per Jones et al., (2021). For each line, ten individual plants per line were embedded and sectioned, and 5-10 root sections were collected per plant, stained using the method described in Atkinson and Wells (2017), and imaged using confocal laser scanning microscopy. Basic fuchsin staining was conducted at a concentration of 0.01% following sectioning. A single image section per plant was selected based on quality and representation, then measured using FIJI (Schindelin et al., 2012). Cells were classified into layers in concentric rings from the endodermis outwards. The diameter of each layer was measured, as was the number of cells in each layer, along with the diameter of the endodermis, number of endodermal cells, and number of cells in the stele. Diameters were measured using the ruler tool. At each layer, diameter was measured from 5 points around the circumference of the layer, measuring the maximum distance between points on the layer, then the mean was taken of these 5 points for each layer. Epidermal cells had poor dye penetration, and a reduced fluorescence on the confocal microscope, and so could not be reliably counted.

### Root removal treatments and imaging

For the root removal experiment, plants were grown in Schenck-Hildebrandt (SH) media. For the control treatment, no manipulation was undertaken. In the root removal treatment, all visible roots were removed from colonies daily using ethanol sterilised surgical scissors. For *Spirodela polyrhiza* and *Lemna minor*, each treatment consisted of five individual flasks, each seeded with 3 colonies onto 100 ml of media. Individual flasks were treated as a replicate and flasks were arranged randomly in the growth cabinet and re-randomized daily. For *Pistia stratiotes*, each flask was seeded with a young individual plant with 3 emerged leaves visible to the naked eye, to a total of 7 plants/treatment. The treatment regimen was conducted for 11 consecutive days.

Plants were imaged daily in their flasks from beneath, utilising a transparent raised platform featuring a water bath in which to place the flasks to correct for the optical distortion. Images were processed using FIJI to measure frond or aerial tissue area. For duckweed flasks, RGB images were split into their constitutive 8-bit channels, and the blue channel retained. Frond tissues alone were then selected using the threshold tool and area measured. For *Pistia*, images were again split, but the red channel retained. This was then subject to gaussian blur (sigma = 7.0) and again only the aerial tissues selected using the threshold tool. In rooted samples where this alone was not sufficient to separate frond and root, the select polygons tool was used to exclude any additional root captured by thresholding.

### Ionomic analysis

Samples were harvested immediately following the root removal experiment. Prior to harvesting, roots were removed from fronds or aerial tissues and washed 3 times for 2 minutes with MilliQ water. Samples were placed in pre-weighed Pyrex test tubes, and dried at 88°C for 24h. Then, dry weight was recorded, and 1 ml concentrated trace metal grade nitric acid Primar Plus (Fisher Chemicals) spiked with in internal standard was added to the samples that were further digested in DigiPREP MS dry block heaters (SCP Science; QMX Laboratories) for 4 hours at 115°C following the method adapted from Danku et al.,2013. After digestion, samples were diluted to 10 mL with 18.2 MΩcm Milli-Q Direct water and elemental analysis was performed using an ICP-MS, PerkinElmer NexION 2000 and twenty-three elements were monitored (Li, B, Na, Mg, P, S, K, Ca, Ti, Cr, Mn, Fe, Co, Ni, Cu, Zn, As, Se, Rb, Sr, Mo, Cd and Pb). To correct for variation within ICP-MS analysis run, liquid reference material was prepared using pooled digested samples, and run after every nine samples. Sample concentrations were calculated using external calibration method within the instrument software. Further data processing including calculation of final elements concentrations was performed in Microsoft Excel.

### Statistical analyses

All statistical analyses were conducted in GraphPad Prism version 9.0 (graphpad.com). For the anatomical dataset, principal coordinates analysis was conducted on 19 variables and 210 rows utilising parallel analysis with 1000 simulations and a random seed. For root removal experiments, two-way repeated measures ANOVA was performed, followed with Sidaks’ multiple comparisons test to establish differences in growth on a per-day basis. For nutrient concentration comparisons generated by ionomic analyses, data were compared with one-way ANOVA followed by Sidak’s multiple comparison’s test to establish differences in concentration between individual nutrients. Log2 fold changes generated from ionomic data were calculated as Log2(elemental conc. roots removed)-Log2(elemental conc. rooted).

## Acknowledgements

We acknowledge the Nottingham Future Food Beacon for support with ionomics, and Walter Lämmler from the Landolt Duckweed Collection and Klaus J Appenroth from the Friedrich Schiller University for kindly supplying material used in this study.

